# Effects of ceratotrichia diameter and packing density on chimaera pectoral fin kinematics

**DOI:** 10.1101/2025.08.11.669704

**Authors:** Duncan W. G. Kennedy, Kayla C. Hall, Cassandra M. Donatelli, Kelsey N. Lucas

**Affiliations:** Biological Sciences, University of Calgary, 2500 University Dr NW, Calgary, AB T2N 1N4, Canada; Department of Biology, University of Washington, 1410 NE Campus Pkwy, Seattle, WA 98195, United States; School of Engineering and Technology, University of Washington Tacoma, 1900 Commerce St, Tacoma, WA 98402, United States

**Keywords:** chimaera, swimming, kinematics, pectoral fin, fin ray

## Abstract

Chimaeras – an ancient group of cartilaginous fishes – swim by flapping their pectoral fins in a distinctive locomotory mode, termed flapping flight, that induces an undulatory wave traveling from the leading edge to the trailing edge of the fin. Recent work on bony fishes has shown that fins with internal structure (fin rays) may behave differently than models made of a single material. Our goal was to understand the potential significance of internal structure for the swimming kinematics of chimaeras. We designed an actuation system based on the kinematic patterns of real chimaeras and examined artificial fins of varying fin ray diameter and packing density to investigate how internal anatomy influences pectoral fin tip amplitude, leading-edge curvature, and the induction of undulatory waves. Both diameter and packing density influenced fin kinematics, but diameter had a much larger effect on fin tip amplitude over the tested ranges. Additionally, established rules for undulatory waves outlined in the flexible foil literature did not hold for our fin models. Our work provides new insights into the anatomical parameters that could influence the evolution of chimaera flapping flight, and pectoral fin locomotion more broadly, and provides direction for possible biomimetic applications.

## Introduction

Fish show remarkable diversity in the form and function of their fins and bodies, supporting a wide range of locomotory modes. In recent years, considerable effort has gone toward understanding the contributions of body and fin material and structural properties to swimming capabilities. These studies have shown that morphology, kinematics, and swimming performance are tightly integrated [1]. Much of this work has relied on flexible foil models with largely homogenous structure, often based on the body and caudal fin [reviewed in 1].

Unlike these models, fish fins are not homogenous. In both bony and cartilaginous fishes, fins are made of relatively stiff fin rays embedded in more flexible tissue [2–6]. There is evidence that fins with supporting fin rays behave differently than those without an internal structure [7]. Here, we take a first principles approach to study how internal fin structure influences pectoral fin kinematics. Through this investigation, we seek to establish a foundation for future research exploring other factors that might impact the kinematics of propulsive surfaces with embedded internal structures.

The significance of internal structure for pectoral fin function is best understood in bony fishes, which actively deform the surface shape and tune the bending stiffness of their fins. This is accomplished not by altering the fin’s intrinsic material properties, but through muscular bending of bilaminar fin rays called lepidotrichia at the base of the fin [2,4]. When this bending occurs, the collagenous membrane between fin rays is stretched until it resists further bending and effectively stiffens the fin [7]. Materials with supporting fin rays behave differently than single material, as evidenced by fish increasing their swimming performance by actively tuning fin stiffness through fin ray curvature [8,9].

Although cartilaginous fishes share this fundamental structure of pectoral fins (rays supporting tissue), the anatomy and control of pectoral fins differs greatly between bony and cartilaginous fishes [2,3,6,10]. Unlike their bony counterparts, cartilaginous fishes are not known to actively tune the stiffness of their fins and vary drastically in pectoral fin anatomy. This results in a gradient of control over fin surface shape between the three major groups: batoids (skates and rays), sharks, and chimaeras.

The fin rays of batoids are segmented into radials attached via thin muscles originating from the base of the pectoral fin [11]. When these muscles are actuated, the pectoral fins deform into sizeable undulatory waves [12–15]. Like batoids, sharks and chimaeras have radials, but they do not possess intricate pectoral fin musculature and are only capable of actuating their pectoral fins at the base [16–18]. Distal to the radials of both sharks and chimaeras are thin filaments of elastic proteins called ceratotrichia which cannot be actively bent [3,5,6,19,20].

Chimaeras propel themselves using a mix of undulation and oscillation, making them an intriguing group of pectoral swimmers. Their unique mode of locomotion, aptly named flapping flight due to the oscillation of enlarged pectoral fins, generates an undulatory wave that travels from the leading edge to the trailing edge [21]. Because chimaeras do not have the anatomy necessary to actively control the curvature of their fin rays and hence the stiffness and surface shape of their fins, their pectoral fins likely deform passively during swimming. However, passive flexibility still shapes swimming biomechanics. For example, passive flexibility can increase swimming performance up to a point [22], after which an increase in flexibility prevents fins from producing coherent kinematics and generating appreciable propulsive forces [7].

Given the tight links between internal structure, kinematics, and performance, the underlying structure of chimaera pectoral fins may have strong implications for their life history and ecology. Particularly, a continuum of manoeuvrability and cruising efficiency exists for paired fin swimmers between undulatory and oscillatory motion [15]. Undulatory movements could aid in manoeuvring around obstacles and hunting prey in cluttered, benthic environments, whereas oscillatory movements might contribute to efficient migration for those species travelling inshore for spawning [15,23–25]. If fin ray structure influences the pectoral fin’s bending kinematics, then where a chimaera falls on the undulatory-oscillatory continuum may be determined in part by internal fin structure. However, the connections between fin ray structure and overall fin kinematics is complex and requires further exploration [1]; it is unclear what specific aspects of fin ray structure are important for determining swimming capabilities.

Here, we focus on fin ray diameter and packing density. If it is assumed that fin rays have a circular cross section, then the bending stiffness of a ray is proportional to the second moment of area, *I*, for a circular rod:

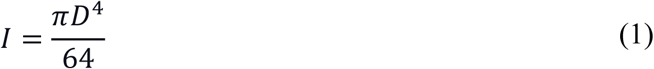

where *D* is the diameter of the rod. Therefore, the bending stiffness of a ray is also proportional to the diameter of the ray. Additionally, stiffness increases with the number of supporting rays within a fin. Given the connections between stiffness and swimming performance [1,8,22], we seek to clarify how fin ray diameter and packing density alter swimming kinematics using flexible artificial fins that mimic the pectoral fins of *Hydrolagus colliei* – a northeastern Pacific chimaera whose range extends from Alaska to Costa Rica [23,26,27]. We chose this species because it is the most readily accessible in museum collections and its swimming is the most well-studied among extant chimaeras [21,26–28]. We predicted that fins with thinner and fewer rays would experience more bending, exhibited by larger fin tip amplitudes, curvature, and induced undulatory waves.

The three primary findings of this study include 1) fin ray diameter and packing density both influenced pectoral fin kinematics; 2) diameter had a much greater effect on fin tip amplitude than packing density over the tested values of each; and 3) established rules in the literature for undulatory waves were not faithfully followed by our flexible fin models. If genetic underpinnings exist for fin ray diameter and packing density in chimaeras, as they do in other fishes [29–31], then our findings suggest that chimaera fin ray structure might have been shaped by the swimming performance demands of chimaera’s environments and behaviours. While we focus on chimaeras, our first principles approach to understanding the impacts of these parameters on kinematics allows for broader translation of our findings to a widespread form of fin locomotion and points to relevant design principles for fin-actuated underwater vehicles.

## Materials and Methods

### Artificial Fin Design

To study the effects of fin ray anatomy on chimaera pectoral fin kinematics, we constructed nine artificial fins of varying fin ray diameters and packing densities. Pectoral fin photographs were taken of five *H. colliei* specimens from the Field Museum of Natural History’s (FMNH) Division of Fishes (Table 1). We selected several fully developed specimens – those with a body length greater than 40 cm – with pectoral fins in good condition [23]. Outlines of the fin and fin muscle were then traced with tpsDig2 (Rohlf, F. J., Stony Brook, NY, USA).

**Table 1.** Specimens from the FMNH used for fin model design.

| Family | Species | Catalogue Number |
| --- | --- | --- |
| Chimaeridae | *Hydrolagus colliei* | 17121 |
| Chimaeridae | *Hydrolagus colliei* | 17123 |
| Chimaeridae | *Hydrolagus colliei* | 117181 |
| Chimaeridae | *Hydrolagus colliei* | 117182 |
| Chimaeridae | *Hydrolagus colliei* | 117183 |

**Table 2.**
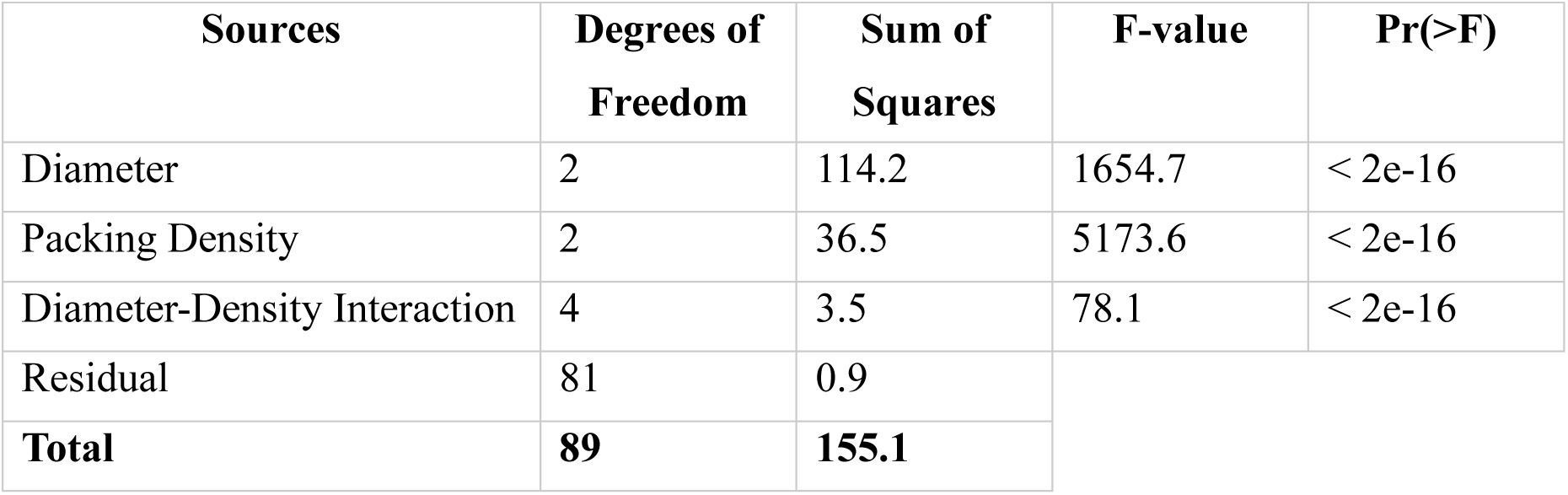
Results of the ANOVA conducted on fin tip amplitude.

This landmarking scheme captures chimaera pectoral fin shape with three fixed landmarks and two sliding semilandmark curves (Figure 1A). Fixed landmarks were placed at readily identifiable homologous points: the anterior fin-muscle connection (point 1), fin tip (point 2), and posterior fin-muscle connection (point 3). One curve outlines the leading edge with 19 sliding semilandmarks and the other outlines the trailing edge with 31. Only every fourth sliding semilandmark was kept; the remaining helper points were culled prior to statistical analysis to ensure appropriate power was retained [32]. The additional helper points assisted in constraining sliding semilandmarks to accurate positions along the curve [33]. The R package *geomorph* (4.2.3; [34,35]) was used to produce an average *H. colliei* fin and fin muscle outline from the five museum specimens. In obtaining a representative fin outline, it was assumed that pectoral fins maintain constant shape between sexes and sizes for fully developed individuals.

**Figure 1:**
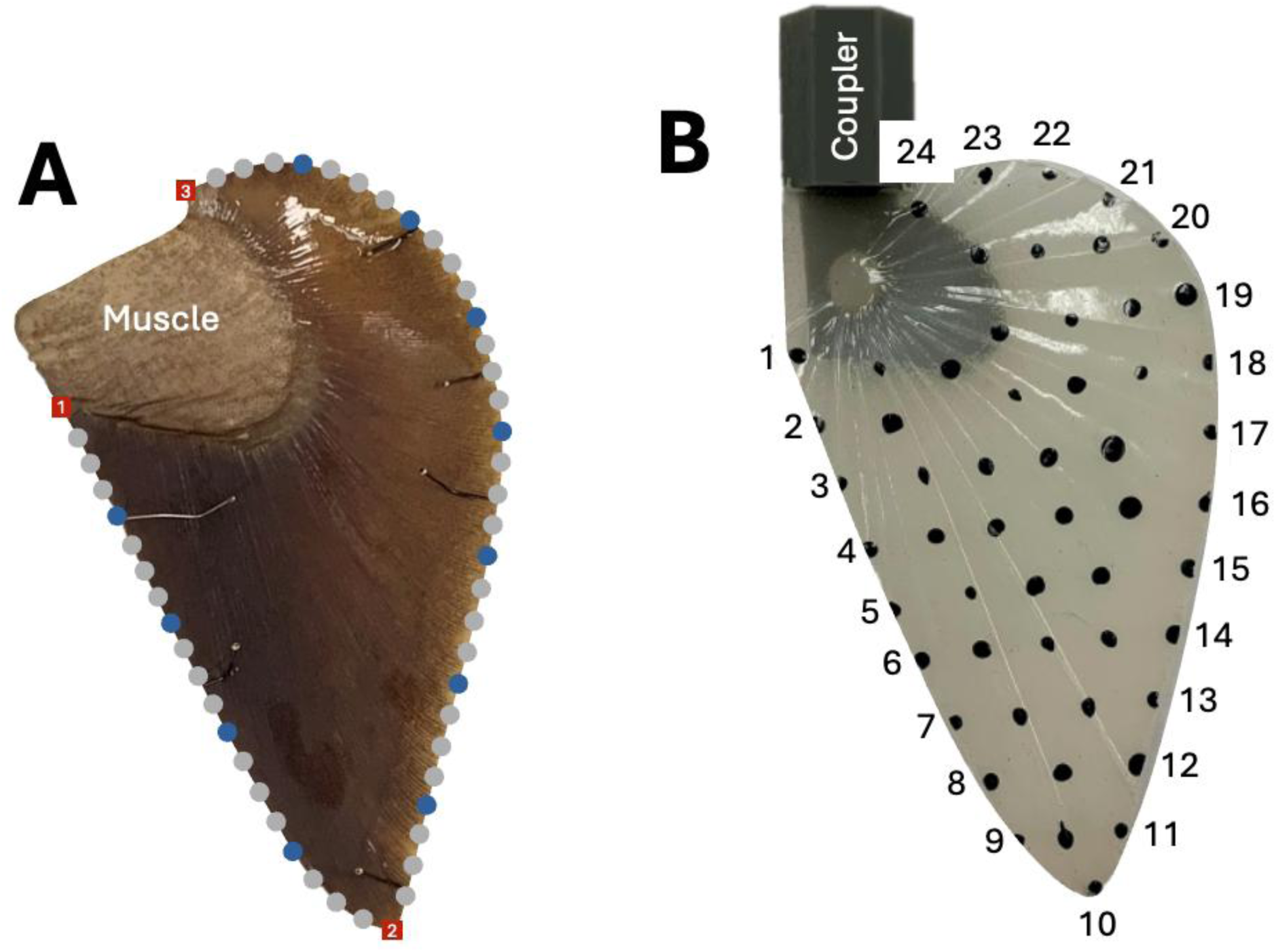
**A)** *Hydrolagus colliei* (FMNH 117183) **pectoral fin.** Muscular region, as labelled in the figure, is used to flap the fin. Red squares indicate fixed landmarks (1: anterior fin-muscle connection, 2: fin tip, 3: posterior fin-muscle connection), blue circles sliding semilandmarks, and grey circles helper points. **B) Example artificial fin fashioned from Ecoflex 00-30.** Every fin has a 3D resin printed fin-robot coupler, as labelled in the figure, which connects the fin to the fin-flapping robot. Thirty pieces of 0.7 mm monofilament fishing line are embedded in this particular fin. Black dots were painted on fins for point tracking purposes. Points 1 through 10 compose the leading edge, and points 10 through 24 compose the trailing edge.

Using the representative fin outline, a fin mold was designed in Fusion 360 (Autodesk, Inc., San Rafael, CA, USA) and printed on a Form 3+ resin printer (Formlabs, Somerville, MA, USA). Monofilament fishing line of varying diameters (0.3, 0.5, and 0.7 mm) and packing densities (14, 30, and 44 strands of fishing line) were threaded through holes in the mold to mimic fin rays (Figure 1B). A fin-robot coupler based on the fin muscle outline, designed in Fusion 360, and printed on the Form 3+ resin printer, was also embedded in the mold. We then filled the mold with Ecoflex 00-30 (Young’s modulus = 0.125 MPa), a flexible silicone rubber with biologically relevant material properties akin to those of zebrafish larva [36]. Because of the different filament embedding schemes, artificial fins follow the naming convention diameter-packing density. For instance, 30 strands of fishing line 0.7 mm in diameter are embedded within the 0.7-30 fin (Figure 1B).

The artificial fins of this study compromise between ease of construction and biological realism. Though chimaera pectoral fins have thinner and far more fin rays than the artificial fins, it is impractical to embed so many rays of such small diameter in artificial fins without resorting to advanced manufacturing methods. Further, this design choice enabled control over fin ray diameter and packing density. This design philosophy has previously been successful for simple physical models exploring the underlying principles of complex fish structures [37–39] and enables exploration of the central question of this study: to what extent do fin ray diameter and packing density influence kinematics.

To track fins during kinematic trials, 54 black dots were painted on the fins approximately 1 cm apart. All fins had a frontal area of 5000 mm², an average thickness of 5 mm, and a leading-edge angle of 26.1⁰. This angle is the average angle between the left-right axis of the fish and the leading edge of the pectoral fin for the five *H. colliei* specimens examined in this study. Angles were measured manually from the photos collected at the FMNH in ImageJ (Rasband, W.S., U. S. National Institutes of Health, Bethesda, MD, USA).

### Robot Design and Verification

We constructed a fin-flapping robot (Figure 2) to “swim” the artificial fins. A NEMA 17 motor flapped the fins, and a 5V stepper motor controlled the pitch of the fins. Both motors repeatedly rotated the fins clockwise then counterclockwise, in phase, producing motion like that of real chimaeras. The robot was housed in a 75.2 cm by 31.7 cm by 43.2 cm tank filled with freshwater.

**Figure 2:**
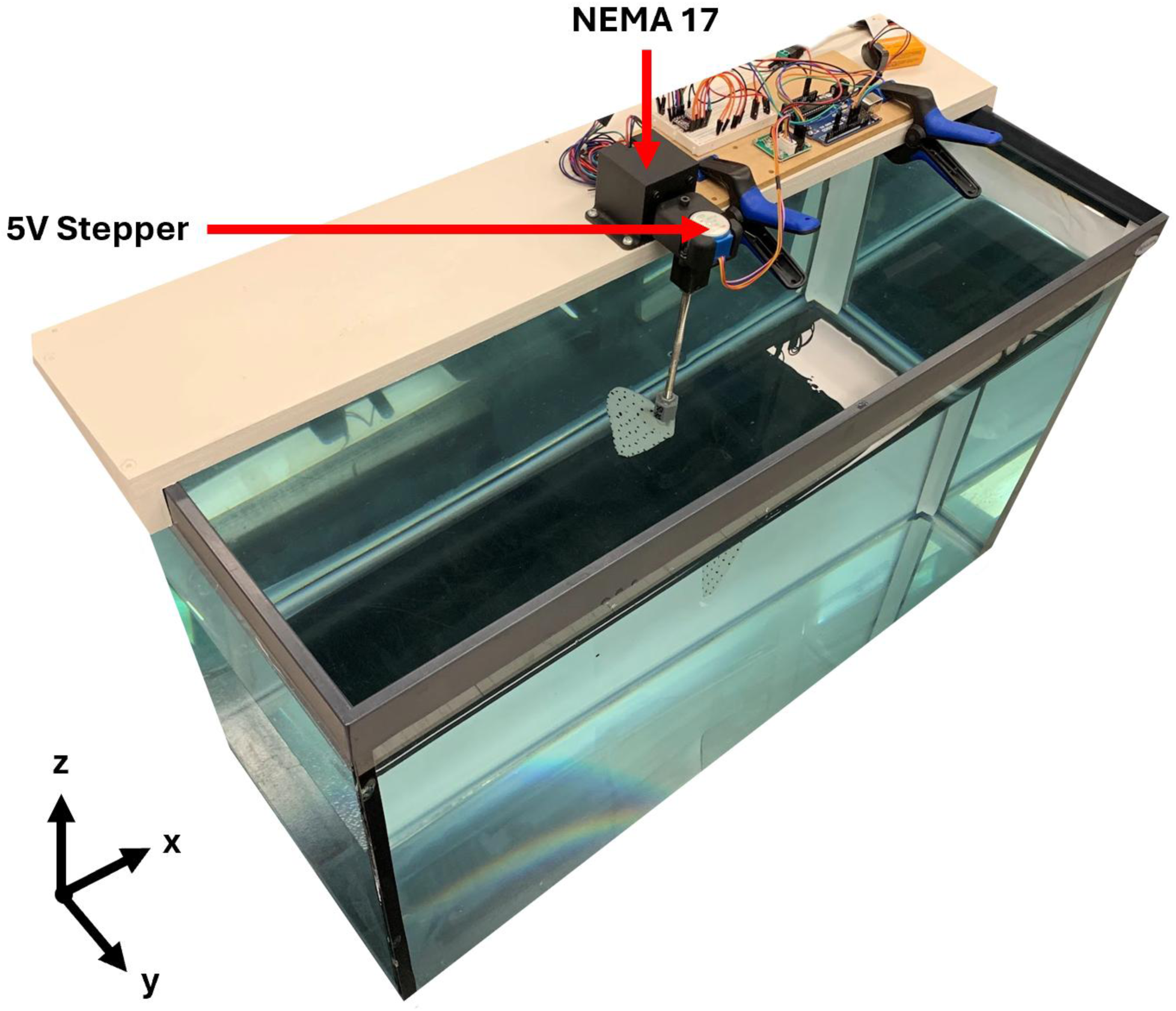
Fin-flapping robot. The NEMA 17 motor flaps the fin around the y-axis. The 5V stepper motor changes the pitch of the fin around the z-axis.

To ensure that the system accurately represents chimaera flapping flight, we programmed the robot with kinematics comparable to those displayed by a live *H. colliei* specimen swimming in a flume with no flow (Figure 3; [27]). Fin beat period (s), fin tip amplitude (FTA, cm), and fin pitch (°) were calculated by tracking and averaging nine pectoral fin strokes of the live specimen using ImageJ. To standardize the position and orientation of the specimen, the base of the dorsal spine was set to the origin and body pitch – the angle between the vector from snout to cloaca and the x-axis – was set to 0⁰. A stroke begins when the fin tip reaches its maximum amplitude (fin tip maximum) at the top of the upstroke, continues while the tip traverses to its minimum amplitude (fin tip minimum) on the downstroke, and ends when the tip regains its maximum amplitude on the upstroke (Figure 4). Fin beat period is the time it takes to complete a stroke.

**Figure 3:**
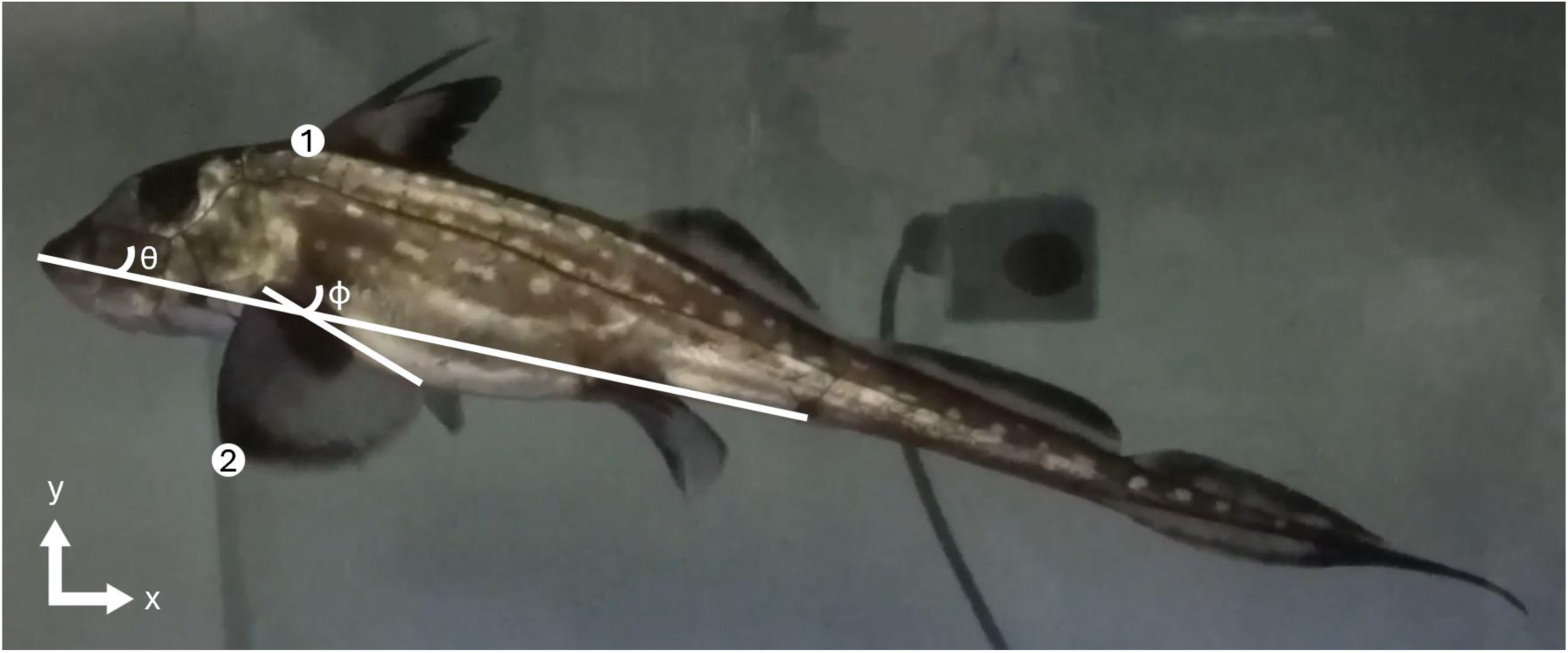
A live *Hydrolagus colliei* specimen swimming in a flume with no flow. [27]. Point 1 marks the base of the dorsal spine, point 2 marks the tip of the pectoral fin, θ is body pitch – the angle between the vector from snout to cloaca and the x-axis, and Φ is fin pitch – the angle between the base of the fin at the bottom of the downstroke and the x-axis. The kinematics of the fin-flapping robot were based on these measurements.

**Figure 4:**
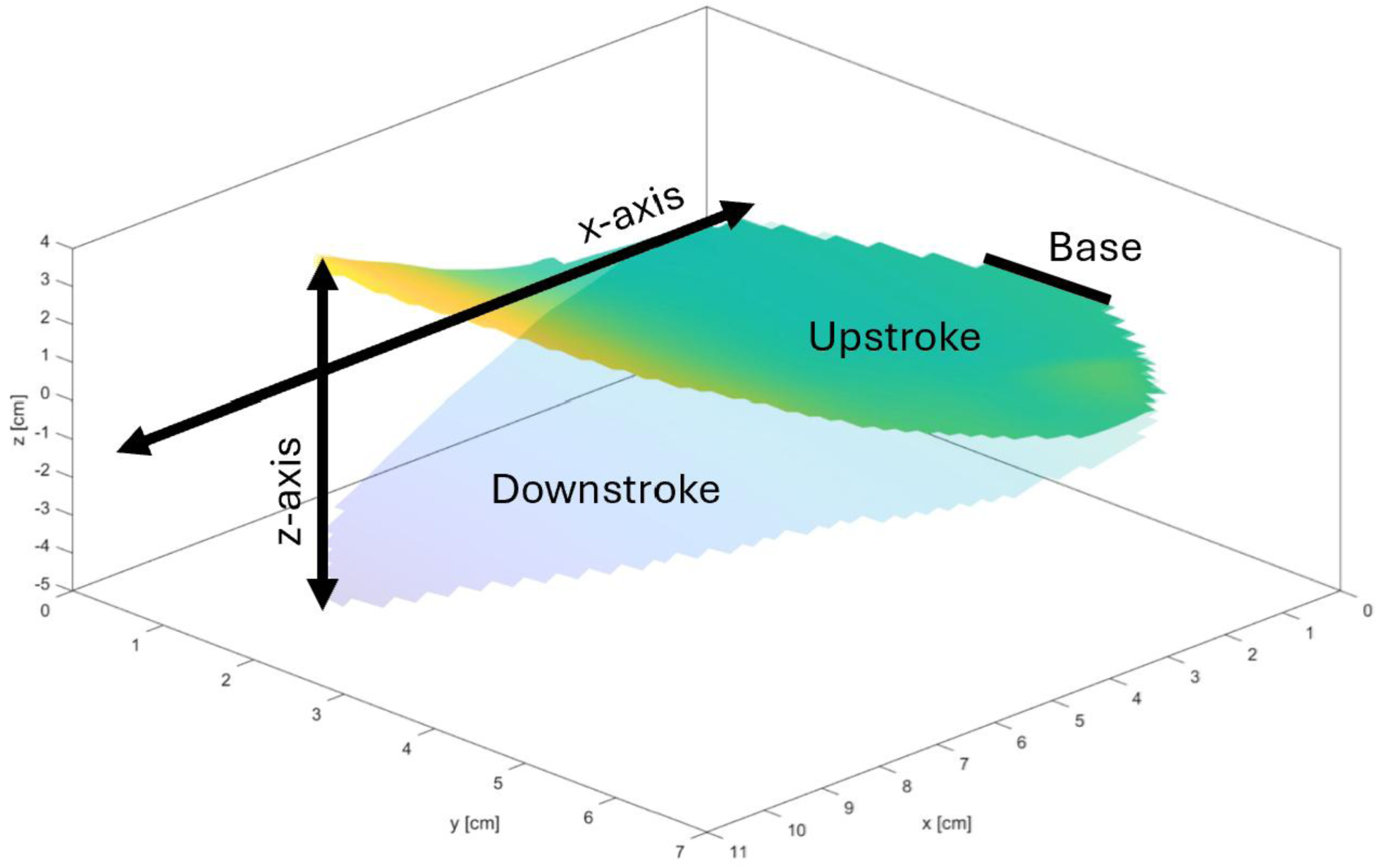
Translation and axis alignment of the artificial fins. The fin base is translated to the same position throughout the stroke. The x-axis is aligned with the leading edge of the fin at midstroke. The z-axis extends from the tip of the fin at the bottom of the downstroke to the tip of the fin at the top of the upstroke. The color gradient represents the amplitude (z-axis value) of each point on the fin, with warmer colors indicating more positive amplitudes and cooler colors indicating more negative amplitudes.

The period of the live specimen in the flume falls within the range of periods exhibited by *H. colliei* found in nature, as verified with four videos recorded in the wild [27]. FTA is the distance between fin tip minimum and maximum. Fin pitch is the angle between the fin base and the x-axis.

Based on these measurements from the live specimen, the robot was programmed to beat the fins with a period of 1.16 s, a NEMA 17 rotation angle of 12.3⁰ (which is equivalent to a linear amplitude between 3.6 and 4.8 cm at the fin base), and a pitch of 15.1⁰. Nuances in pectoral fin kinematics would not have been obvious with the relatively small FTA the live specimen exhibited; therefore, we substantially increased the robot’s FTA. All other robot parameters were identical to those displayed by the live specimen.

### Kinematic Data Collection

We filmed each of the nine artificial fins at 60 fps with two GoPro Hero10s (GoPro, Inc., San Mateo, CA, USA) and an Apple iPhone XR (Apple, Inc., Cupertino, CA, USA). One Hero10 filmed the fins from the top left corner of the tank, the other Hero10 filmed from the bottom right corner, and the iPhone XR filmed from the centre; camera angles were similar to those recommended by and shown in Figure 3B of Theriault et al. [40]. Filming order of the fins was randomized.

To reconstruct the 3D movement of the fins and calibrate the cameras, we waved a calibration wand through all three camera’s shared field of view. Both ends of the wand were tracked through 86 shared frames using the point-tracking software DLTdv8 [41]. The calibration software easyWand5 computed relative camera positions and orientations using the wand points and six stationary background points [40]. Our measures of calibration inaccuracy were 0.003807 for the standard deviation of the estimated wand length divided by its mean and 0.50, 0.51, and 0.47 pixels for the root mean squared distances between the original and reprojected points, indicating high accuracy in the calibration of our cameras [40].

Filming began after the robot ran for a minimum of one minute to ensure that the system reached steady-state – when the same flow is established around the fin for every stroke. A flashlight was turned on and off in the cameras’ shared field of view for synchronization purposes. Ten strokes were then recorded for each fin. Videos were imported into DLTdv8, synchronized with the flashlight strobe, and the 54 black dots on each of the nine artificial fins were tracked through all ten strokes.

### Kinematic Analysis

Because chimaeras cannot translate the base of their pectoral fins while swimming, we mathematically removed translation at the base of each artificial fin using MATLAB (MathWorks, Inc., Natick, MA, USA). Point 1 on each artificial fin, which corresponds to the base of the fin in live chimaeras, was set at the origin by subtracting the amplitude of point 1 from the amplitudes of all 54 points. Fin orientation was standardized by aligning the vector from fin tip minimum to fin tip maximum with the z-axis and by aligning the leading edge of the fin at midstroke with the x-axis using Rodrigues’ rotation formula (Figure 4).

FTA was measured over ten strokes and analyzed using ANOVA. A post-hoc orthogonal polynomial contrast was performed to disentangle linear and non-linear effects of fin ray diameter and fin ray packing density on, and their relative contributions to, FTA [42].

To calculate leading-edge curvature, we first fit a second-degree polynomial to the leading edge of each artificial fin using MATLAB’s “polyfit” function. A second-degree polynomial was selected as it is the lowest order polynomial to accurately outline the leading edge. Overfitting becomes more likely as polynomial order increases [43]. The curvature, κ, at a point *i* along the leading edge is approximated with:

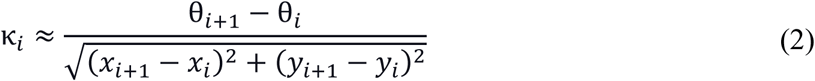

where θ is the angle between the vector from point *i* to point *i*+1 and the positive x-axis, and *x* and *y* are point coordinates (Figure 5).

**Figure 5:**
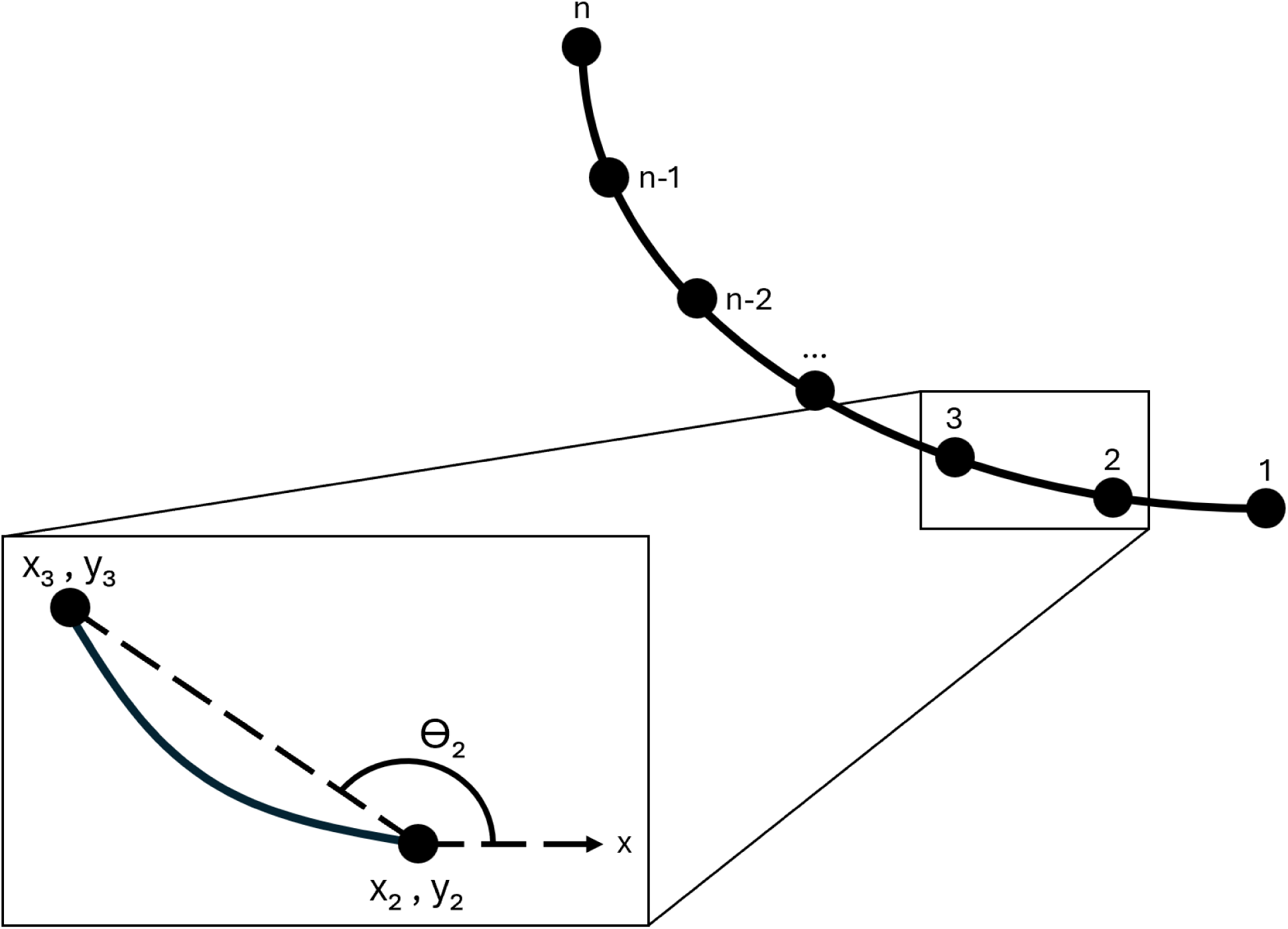
Calculating curvature. A polynomial is fit to either the leading or trailing edge of the artificial fin. The polynomial is then split into several smaller arcs bounded by points. The equation for the curvature, *K*, of an arc requires 1) the coordinates, (*x, y*), of the arc’s bounding points and 2) the angle, θ, between the vector connecting the bounding points and the positive x-axis.

Several of the artificial fins generated an undulatory wave that traveled distally along their trailing edge. The presence of a wave is indicated by an inflection point – the point where curvature changes sign. Undulatory wave curvature is calculated in the same way as leading-edge curvature (Equation 2), but in this case with a fourth order polynomial instead of a second order polynomial. We calculated wave interval – when a wave begins and ends – by recording when an inflection point appears and subsequently disappears. Intervals have been normalized to the wave period and are reported as percents. Average wave speed (m/s) was calculated by tracking and averaging the position of an inflection point through time [44].

### Data and Code Availability

Kinematics videos and code will be uploaded to a publicly available repository (e.g., Dryad, GitHub, etc.) upon manuscript acceptance, and this text will be updated accordingly.

## Results

Artificial fins shared various kinematic patterns as they were flapped by the robot. Wave amplitude increased from base to tip, undulatory waves appeared on the trailing edge of several fins and travelled distally, and the fin tip lagged slightly behind the fin base until stroke reversal when the tip would rapidly accelerate past the base.

### Fin Tip Amplitude and Leading-Edge Curvature

FTA was predominantly governed by the linear effects of fin ray diameter and fin ray packing density. FTA decreased 30% (from 8.6 to 6.0 cm) and 19% (from 7.8 to 6.3 cm) over the ranges of diameter and packing density, respectively. The linear effects of diameter accounted for 63.9% of the variation in FTA, while the linear effects of packing density accounted for 23.1% (Table 3). Diameter played a much larger role in dictating FTA.

**Table 3.** Results of the orthogonal polynomial contrast conducted on fin tip amplitude. Percent sum of squares describes the relative effect size of each source (diameter, packing density, etc.) on the variation seen in amplitude. Contributions from diameter and packing density are further partitioned into contrasts describing linearity and deviation from linearity.

| Sources | Degrees of Freedom | Sum of Squares | Percent Sum of Squares |
| --- | --- | --- | --- |
| Diameter | 2 | 114.2 | 73.6 |
| Linear | 1 | 99.1 | 63.9 |
| Deviation | 1 | 15.1 | 9.7 |
| Packing Density | 2 | 36.5 | 23.6 |
| Linear | 1 | 35.8 | 23.1 |
| Deviation | 1 | 0.7 | 0.5 |
| Diameter-Density Interaction | 4 | 3.5 | 2.2 |
| Residual | 81 | 0.9 | 0.6 |
| **Total** | **89** | **155.1** | **100** |

The non-linear effects of both diameter (9.7%) and packing density (0.5%) accounted for much less than the linear effects. In Figure 6, each line connecting the average FTA between artificial fins of the same packing density and different diameters bends upwards (the slope becomes less negative). No such trend occurs for fins of the same diameter and different packing densities; the line for the 0.3 mm fins bends downwards, the line for the 0.5 mm fins is straight, and the line for the 0.7 mm fins bends upwards.

**Figure 6:**
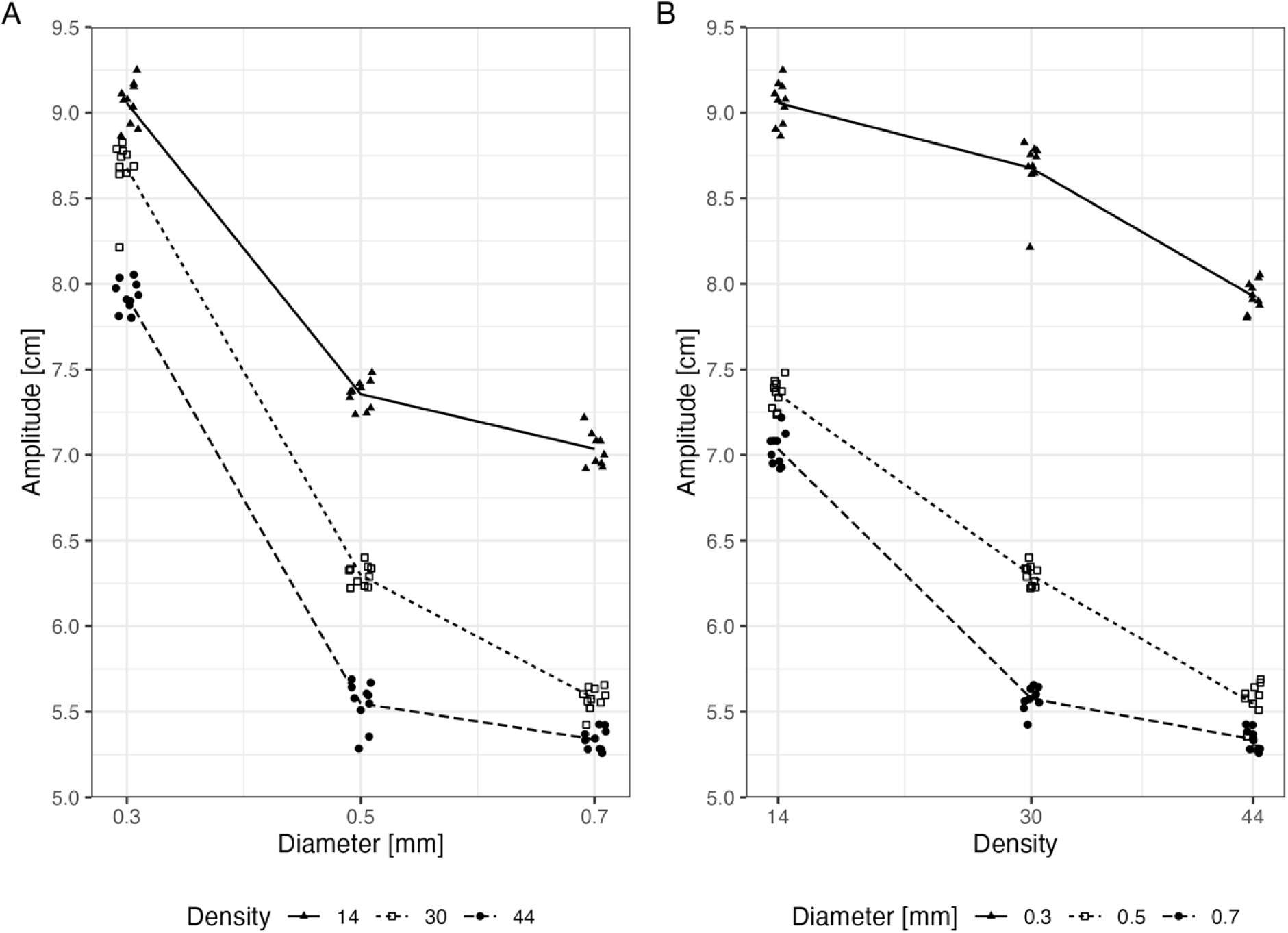
Fin tip amplitude changed with fin ray structure. A) Fin tip amplitude decreased as fin ray diameter increased – shown by the lines connecting the mean amplitudes of artificial fins with the same fin ray packing density. B) Fin tip amplitude decreased as fin ray packing density increased – shown by the lines connecting the mean amplitudes of artificial fins with the same fin ray diameter. Fin tip amplitude is the distance travelled by the tip of an artificial fin and is measured over ten strokes for each fin. Packing density is the number of fin rays embedded in a fin.

The interaction between diameter and packing density had a significant effect on FTA (ANOVA: significant two-way interaction term, p-value < 2e-16, F-value = 78.1, DF = 4). However, it only accounted for 2.2% of the total variation in FTA and thus is negligible when compared to the independent effects of diameter and packing density. Figure 6 displays this finding graphically. Put differently, diameter did not sizeably change the effects of packing density on amplitude and vice versa. Lastly, residuals described only 6.6% of the variation in FTA, an indication that samples are tightly grouped around the mean.

Like FTA, the leading edge-curvature of the fin decreased as diameter, packing density, and distance from the base of the fin increased (Figure 7).

**Figure 7:**
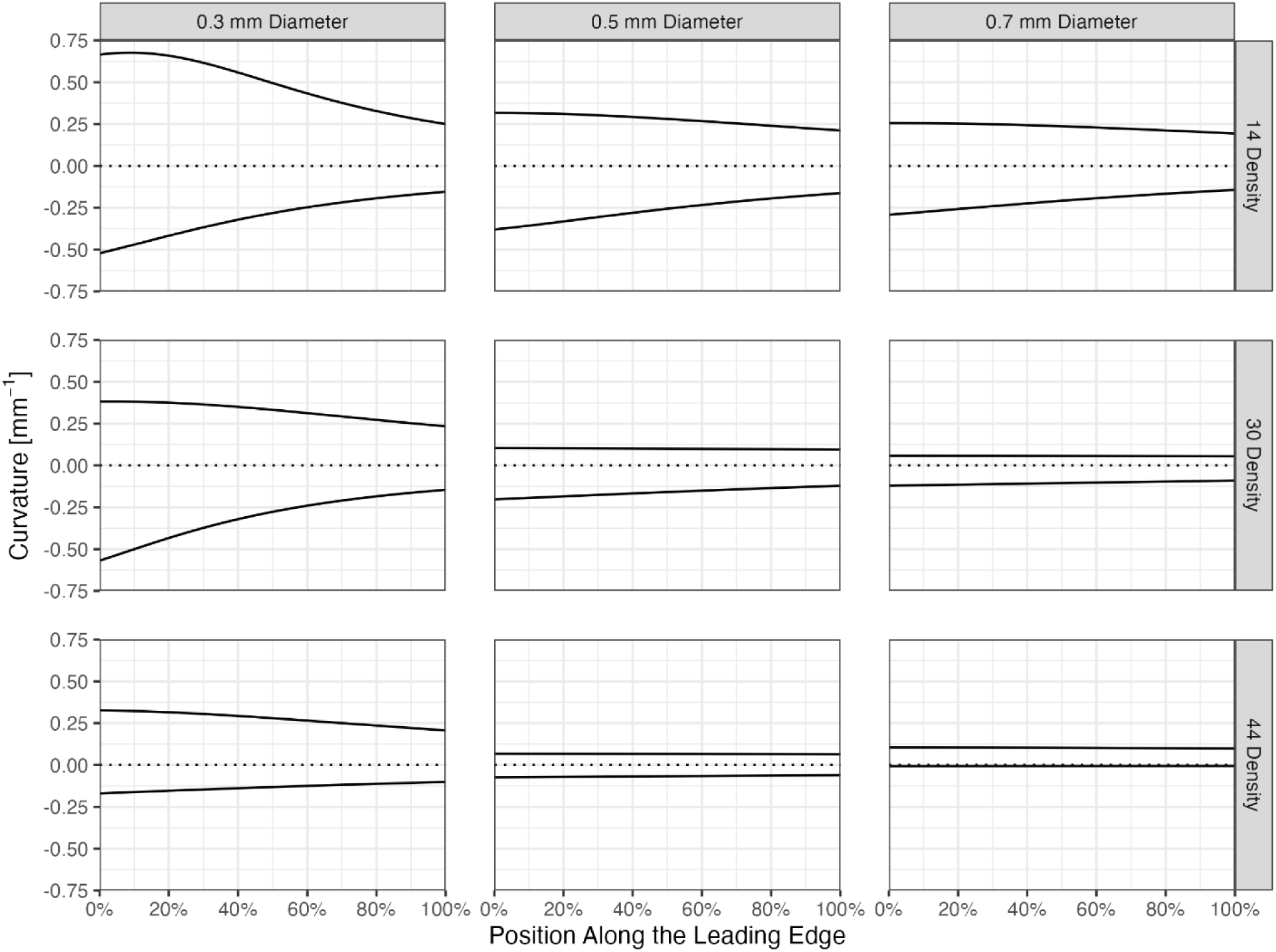
Leading-edge curvature decreased as diameter, packing density, and position increased. The solid line above the dotted x-axis in each plot is the curvature of the leading edge of the fin at the top of the upstroke. The line below the x-axis is the curvature at the bottom of the downstroke. These two lines bound the entire range of curvature an artificial fin will exhibit throughout a stroke.

### Trailing Edge Undulatory Wave

There was no discernable pattern, based on diameter and packing density, for the presence of an undulatory wave travelling distally on the fin’s trailing edge (Figure 8). The fins that did exhibit a wave (0.3-14, 0.3-44, 0.5-14, 0.5-30) did so near the end of the upstroke. The wave on the 0.3-44 and 0.5-14 fins continued through stroke reversal and ended on the downstroke.

**Figure 8:**
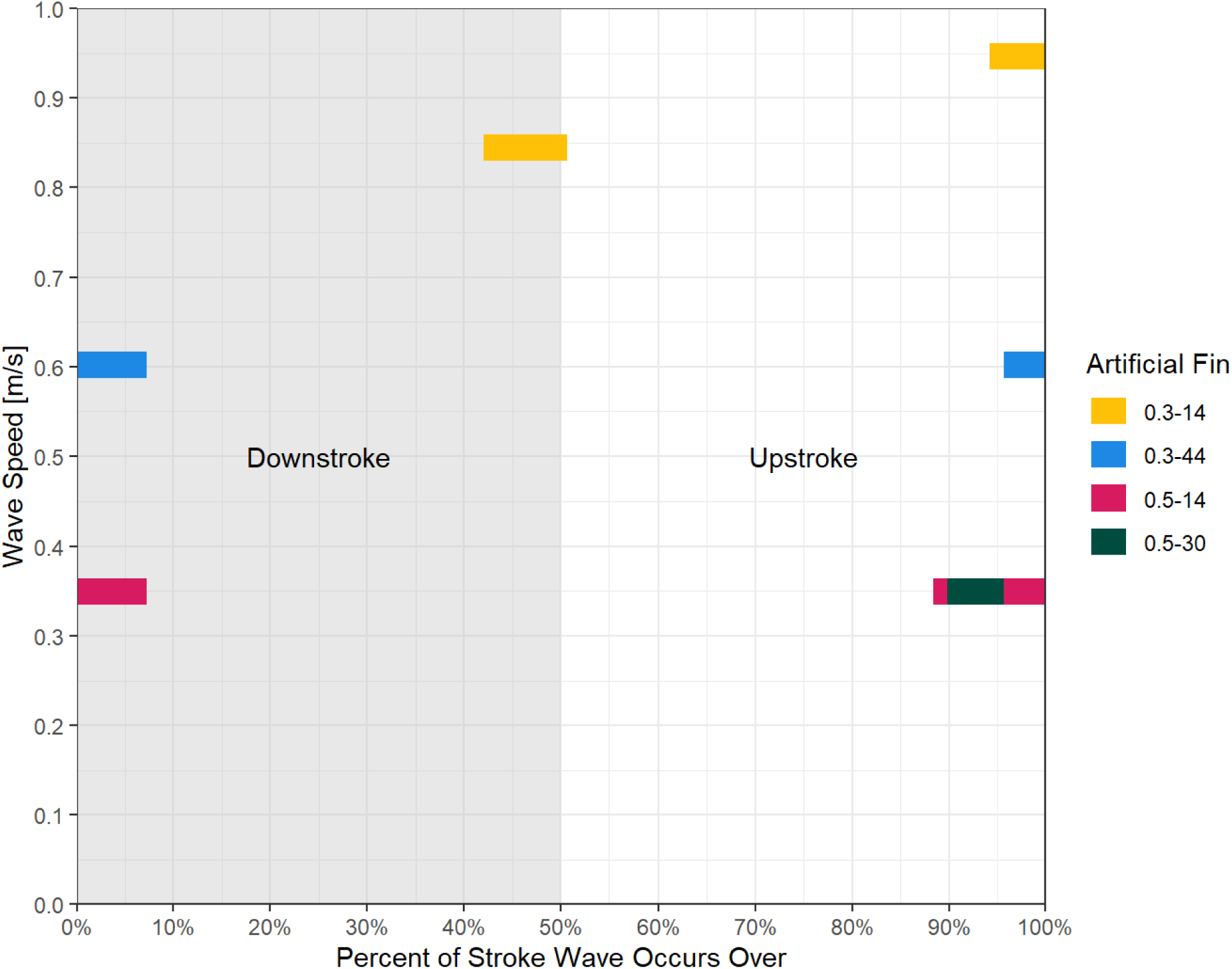
Only four of the nine artificial fins generated an undulatory wave. The 0.3-14 fin was the only fin that generated two waves: one during the downstroke and another during the upstroke. The three other artificial fins only generated one wave near the end of the upstroke which continued through stroke reversal and ended during the downstroke. Strokes are continuous; therefore, colored bars wrap around the x-axis to the other side of the plot (from 100% to 0%). The downstroke occurs from 0% to 50% of the stroke, and the upstroke from 50% to 100%. The y-axis indicates the average speed of the wave as it travels along the trailing edge from fin base to fin tip.

Additionally, the most flexible fin (0.3-14) was the only fin to produce another undulatory wave on its trailing edge during the downstroke. Of the fins that generated a wave, more flexible fins with slimmer and less densely packed rays – such as the 0.3-14 and 0.3-44 fins – tended to have greater average wave speeds. The two fins with rays 0.5 mm in diameter (0.5-14 and 0.5-30) had identical wave speeds (0.35 m/s).

## Discussion

We constructed artificial fins of nine different fin ray diameter and fin ray packing density combinations and actuated these fins with a custom-built fin-flapping robot designed to mimic chimaera flapping flight. An increase in both ray diameter and packing density led to changes in fin kinematics. We hypothesized that fins with thinner and fewer rays would display larger amplitudes at their tips, greater curvature along their leading edges, and undulatory waves because of their flexibility. FTA and leading-edge curvature followed this convention, with flexible fins displaying a higher FTA and leading-edge curvature. Undulatory waves did not: the presences and size of waves did not increase with a reduction in diameter or density.

Research on flapping flight kinematics in chimaeras has been focused on one species, *H. colliei*, and is limited to a few studies [21,26–28]. This study builds on previous research, is the first to explore how fin ray diameter and packing density alter chimaera pectoral fin kinematics and identifies parameters for further exploration in future work on flapping flight.

### Fin Kinematics Depend on Stiffness

An increase in fin stiffness – due to an increase in fin ray diameter and/or packing density – led to a decrease in FTA. Individual rays act as cantilever beams with a load applied at their free end (along the distal border of the fin) due to hydrodynamic and inertial forces during flapping flight [11]. The deflection of a cantilever beam’s free end is inversely proportional to second moment of area, and thus diameter (Equation 1). As diameter increases, the deflection of a ray decreases, and FTA decreases. Tangorra et al. [22] designed a biorobotic pectoral fin modelled after the pectoral fins of bluegill sunfish and investigated how fin rays of varying cross sections and stiffnesses affected swimming performance. Rays of uniform rectangular cross section bend considerably at the base but deflect very little along their remaining length, whereas rays that taper in a fashion more biologically faithful to sunfish fin rays deflect along their entire length. Thick fin rays, either those with large diameters (this study) or uniform cross sections, have relatively small amplitudes at their tips. Thin fin rays, those with small diameters (this study) or tapered cross sections, have relatively large amplitudes [22].

Additionally, fin rays do not act independently of one another; the stiffness of each ray contributes to the overall stiffness of the fin [7]. A fin with more rays, including the artificial fins of this study, has more support and a smaller FTA. Nguyen et al. [7] developed a mathematical model of a rayed fin to determine how fishes modulate fin stiffness and discovered that fin ray bending and fin membrane stretching are coupled. When rays bend, the membrane between them is stretched and elongated. Further ray bending results in additional resistance to stretching by the membrane and a stiffening of the fin. The properties of this coupled system change as fin ray packing density increases: a greater number of rays leads to an increase in fine-scale control over the membrane’s surface and in the overall bending stiffness of the fin. However, Puri et al. [45] found that the bending stiffness of zebrafish caudal fins was unaffected when the fin membrane was cut, suggesting that fin stiffness primarily relies on fin rays and not stretching of the fin membrane. Although this appears to be in direct disagreement with Nguyen et al. [7], Puri et al. [45] agree that the material properties of the interray membrane must be considered when examining fin locomotion. Considering these findings, we find it convincing that both the rays and membrane of the artificial fins in this study contributed to the overall stiffness of the fins. The membrane may also stretch and stiffen as the fin and internal rays passively deform through the water, contributing to the non-linearity seen between fin ray packing density and FTA. Future work on artificial and dissected chimaera pectoral fins could examine whether these fishes employ a bending-stretching mechanism during flapping flight with an experimental setup similar to the setup described in Puri et al. [45].

Leading-edge curvature is brought about by a bending of the fin as it travels through fluid; therefore, as fin ray diameter and packing density increase, fins are more resistant to bending, and curvature decreases. Additionally, curvature decreases non-linearly from fin base to fin tip (Figure 7). Inherent structural properties, such as changes in stiffness, can lead to variation in curvature [12,46]. The base is supported by a relatively stiff fin-robot coupler (3D printed resin) which abruptly ends as the main, flexible body (silicone rubber) of the artificial fin begins. This sudden change in material could give rise to the high initial curvature seen between the stiff fin-robot coupler and the flexible fin body; maximum curvature looks to occur at the point of material transition between the stiff foil body and flexible trailing edge in Cleaver et al. [47]. The continuous silicone rubber body of the artificial fins then dampens curvature gradually from base to tip. Though kinematics is not necessarily a predictor of swimming performance [1,48], our data suggest some theoretical future avenues to explore. For example, boundary layer separation might occur if curvature is too high, leading to an increase in drag and a decrease in performance [49]. Conversely, an increase in curvature could strengthen spanwise flow from fin base to fin tip, stabilizing vortices and increasing efficiency [50].

Fin ray diameter played a much larger role in dictating FTA than fin ray packing density. Although this finding might apply to other kinematic measurements, we note that our current findings regarding FTA should not be extrapolated to leading-edge curvature or undulatory wave speed. Dynamic oscillatory tests, following the recommendations of Jimenez et al. [1], could be used to measure the stiffness of artificial fins and disentangle the effects of diameter and packing density on stiffness. If diameter has a larger effect over stiffness than packing density, as it did for FTA, then it would be reasonable to posit that diameter also affects other kinematic parameters to a greater extent than packing density.

We also suggest that changes in diameter may matter more for kinematics specifically in fins with small fin ray diameters. We observed a much larger decrease in FTA between the 0.3 and 0.5 mm models (∼1.5 to 2.5 cm reduction) than the 0.5 and 0.7 mm models (∼0.25 to 0.5 cm reduction). We estimate that the largest chimaera ceratotrichia are ∼0.1 mm in diameter based on photographs of chimaera pectoral fins [26,51]. This is within the diameter range of 0.06 to 0.2 mm reported for elasmobranchs [5], though we do note that many ceratotrichia within a chimaera fin are likely smaller than these maximums and could not be accurately measured from photographs available in the literature. That said, our smallest ray models do appear to reflect the larger end of the biological range, and the range of diameters among our fishing lines (0.4 mm) is larger than the range in diameters among shark ceratotrichia (0.14 mm) [5]. So, while a wider range of models would be required to fully characterize this relationship, our findings suggest that small changes to diameter among chimaera species may be important for their placement on the undulatory-oscillatory continuum that separates species specialized for maneouvrability from those specialized for cruising efficiency [15].

### Flexible Fins Defy Kinematic Paradigms

Unlike FTA and leading-edge curvature, which decreased with an increase in fin diameter, the presence of an undulatory wave on a fin’s trailing edge could not be predicted based on diameter and packing density. The most flexible fin (0.3-14) produced two waves: one on the upstroke and one on the downstroke. Stiffer fins only produced a wave on the upstroke. This finding aligns with our hypothesis that flexible fins would display undulatory waves more readily than stiff fins, as they are less resistant to deforming and bending as they move through the water.

The two stiffest artificial fins that induced undulatory waves on their trailing edges (0.5-14 and 0.5-30) exhibited identical wave speeds. This suggests that as fins become stiffer their kinematics become more similar. A comparable trend appears in FTA, which decreased but began to level off as fin ray diameter increased (Figure 6A). A certain amount of flexibility might be necessary for fins to respond to hydrodynamic forces they encounter and create in the water; fin surface deformations will not occur if a fin is too rigid. Future work involving artificial fins that cover a greater and more finely incremented range of ray diameters and packing densities might indicate that kinematic differences between fins begin to disappear as stiffness increases.

The faster average wave speed of more flexible fins contradicts established rules in the literature [21,46]. According to these rules, changes in stiffness can control propulsive wave speed and stiffer fins generate faster undulatory waves. However, Combes and Daniel [21] found that when flexible fins are flapped at higher frequencies, established performance rules break down. They used a mathematical fluid flow model to determine the relative performance of flexible fins with different shapes and found that at high flapping frequencies, fins with a broad shape can outperform slender fins in terms of both thrust generation and efficiency. That said, these same shapes perform poorly when flapped at low frequencies or when made with stiffer material. Similarly, our findings suggest that kinematic paradigms, such as the effect of stiffness on wave speed, might fall apart when flexible fins are flapped at specific frequencies.

### Possibilities for Fin Ray Evolution

Genes have been shown to regulate fin ray diameter and packing density. Ray packing density is heritable in the guppy and carp [31], and genetic underpinnings for packing density exist in the little skate, a fish closely related to chimaeras [30]. Additionally, the rays of mutant medaka, or Japanese rice fish, do not normally elongate or thicken [29]; suggesting a level of genetic influence over ray diameter in fishes. If similar genetic foundations for diameter and packing density occur in chimaeras, selection pressures might act on these traits to evolve well-adapted pectoral fins with stiffnesses specialized to the environmental niches chimaeras occupy and behaviours they display [7].

Notably, the stiffness of skeletons in batoids affects swimming kinematics, swimming performance, and location in the water column [11,15]. Batoid oscillators typically have stiff, highly calcified skeletons; are more efficient during cruising; and occupy the pelagic zone.

Conversely, batoid undulators have relatively flexible skeletons, specialize in manoeuvrability, and inhabit more benthic environments. Variation in the pectoral fin stiffness of chimaeras could also lead to differences in kinematics, performance, and habitat preferences like those stated above for skates and rays. Although chimaeras likely cannot actively control the stiffness of their pectoral fins, the range of stiffnesses they do express may have been shaped by their benthic lifestyle and unique ecology.

For instance, if an increase in fin ray diameter improves swimming performance, then thick rays may be selected for. However, we would expect ray diameter to increase only until it no longer has a strong influence on kinematics or swimming performance for two reasons. First, the FTA decrease levels off as ray diameter increases. An increase in diameter after this cutoff might have little or no effect on kinematics and swimming performance and thus would not be selected for. Second, if developing and maintaining fins rays is more energetically expensive than developing and maintaining the surrounding pectoral fin membrane, then stabilizing selection might act to limit the thickness of fin rays.

In summary, both fin ray diameter and packing density influence fin kinematics [7,11,22]. By examining the kinematics of artificial chimaera pectoral fins of varying stiffness, we found that both an increase in diameter and packing density led to a decrease in FTA and leading-edge curvature, diameter played a larger role in dictating FTA than packing density, and trailing edge undulatory waves unexpectedly travelled faster on flexible fins. Because genes have been shown to regulate fin ray diameter and packing density in other fishes [29–31], chimaera fin rays may have been shaped by selection pressures related to swimming performance. This first principles study lays the groundwork for future research on chimaera pectoral fin kinematics in relation to stiffness; reveals biologically relevant parameters of interest for further investigation; and could lead to the design of novel underwater vehicle propulsors with optimally tuned embedded fin rays tuned to optimal diameters and packing densities.

## Acknowledgements

We thank Dr. Peter Blenis for his invaluable statistical expertise; the staff and associates at the Field Museum of Natural History’s Division of Fishes for access to and assistance with their fishes collection; Tal Perevolotsky for her help designing and constructing the robot prototype; Drs. Mark Webster, Matthew Kolmann, and Katelyn Mika for sharing their knowledge on geometric morphometrics and fin ray genetics; Camryn Charriere for feedback on an earlier version of the manuscript; and Heather Jamniczky, Adam Summers, and members of the Lucas and Jamniczky labs for their insight.

## Authorship Credits

Conceptualization – DK, KH, CMD, KNL; Data curation – DK; Formal analysis – DK; Funding acquisition – KNL; Investigation – DK; Methodology – DK, KH, CMD, KNL; Project administration – DK, KNL; Resources – KNL; Software – DK; Supervision – KNL; Validation – DK; Visualization – DK; Writing – original draft – DK; Writing – review & editing – DK, KH, CMD, KNL

